# Intellectual ability and cortical homotopy development in children and adolescents

**DOI:** 10.1101/2025.03.24.645014

**Authors:** Li-Zhen Chen, Xi-Nian Zuo

## Abstract

Functional homotopy, defined as the similarity between the corresponding regions of the two hemispheres, is a critical feature of interhemispheric communication and cognitive integration. Throughout development, the brain transitions from broadly connected networks in early childhood to more specialized configurations in adolescence, accompanied by increased hemispheric differentiation and integration. Using longitudinal data and a novel metric of functional homotopy, homotopic functional affinity (HFA), we investigated the developmental patterns of functional homotopy and its relationship with intelligence. Our findings indicate a significant decrease in HFA with age, particularly in higher-order association networks. In addition, adolescents demonstrate stronger, predominantly negative correlations between HFA and intelligence, in contrast to younger children. In particular, individuals with superior intellectual ability experience accelerated decreases in HFA, indicating greater neural efficiency based on advanced hemispheric specialization and differentiation. These findings provide evidence of the neural mechanisms that underlie cognitive development, emphasizing the dynamic interaction between hemispheric organization and intelligence. Our work may have implications for the design of customized educational/clinical interventions to optimize individual developmental strategies.

**Highlights:**

- Substantial decreases in functional homotopy along the unimodal-transmodal axis observed from childhood to adolescence, with associative areas experiencing a more pronounced decrease.
- Dynamic shifts in the correlation between functional homotopy and intelligence in developmental stages.
- Individuals with a higher IQ demonstrate a significantly faster development of hemispheric specialization or differentiation.
- The distinct patterns in the development of functional homotopy in different IQ- groups underscore the complex interaction between brain maturation and intelligence.

## 1. Introduction

The human brain is organized into a complex network of specialized regions, with dynamic interhemispheric communication that enables a wide range of cognitive functions. A key concept in understanding brain organization is functional homotopy, which refers to the functional similarity between mirror regions of the left and right hemispheres. This homotopy supports effective interhemispheric communication, enabling a spectrum of tasks, from basic sensory processes to complex processes (Ocklenburg and Guo 2024). Traditional theories of functional lateralization have dominated our understanding of hemispheric specialization, asserting that certain cognitive functions are primarily managed by one hemisphere. For example, the left prefrontal cortex is commonly associated with language processing and executive functions, while the right parietal cortex is key to spatial and visuospatial processing (Witelson 1976; Bradshaw and Nettleton 1981; Gotts et al. 2013; Friederici 2011). However, these lateralized functions are not fixed or absolute; rather, they are dynamic and context-dependent. Both hemispheres work together to support higher-order cognitive functions, and the interaction between hemispheric specialization and interhemispheric communication plays a crucial role in intelligence (Miller and Cohen 2001; Karnath and Rorden 2012; Frith and Frith 2012). Functional homotopy, in which homotopic regions in each hemisphere perform complementary roles, allows flexible integration of specialized functions, forming the foundation for complex cognitive abilities.

Recent intelligence models suggest that it arises not from isolated brain regions, but from the coordinated activity of distributed brain networks (Colom et al. 2006). The Parieto-Frontal Integration Theory (P-FIT), for example, posits that intelligence emerges from the collaborative functioning of the prefrontal and parietal regions, which are crucial for high-level cognitive processing (Jung and Haier 2007). Furthermore, neural efficiency theories propose that highly intelligent individuals demonstrate more efficient integration of brain networks, allowing them to complete complex tasks with fewer neural resources (Van Den Heuvel et al. 2009). The dynamic network reorganization theory further suggests that intelligent individuals exhibit greater flexibility in adapting their brain networks to meet changing cognitive demands (Braun et al. 2015). These perspectives emphasize that intelligence is rooted in the way the brain integrates and reorganizes its networks, a capacity strongly linked to functional homotopy between hemispheres.

As the brain develops, its functional organization undergoes significant changes. In early childhood, brain networks are broadly distributed, with high levels of interhemispheric connectivity (Zuo et al. 2010; Fair et al. 2009). Over time, there is a shift towards greater functional specialization, particularly along the spectrum from primary sensory regions to higher-order cortical areas (Dong et al. 2021; Sydnor et al. 2023). This shift reflects the brain’s transition from undifferentiated networks to more specialized configurations that support increasingly complex cognitive functions (Buckner and Krienen 2013; Hill et al. 2010; Sydnor et al. 2021). Importantly, this developmental process occurs alongside stronger hemispheric differentiation(Zuo et al. 2010). As specialized functions emerge within each hemisphere, effective inter- hemispheric coordination becomes increasingly important, facilitating the efficient communication and integration of these specialized functions. Consequently, the trajectory of hemispheric differentiation and integration is fundamental to understanding individual differences in intelligence. Studies have shown that people with higher intelligence tend to exhibit accelerated brain development, achieving more efficient brain organization early than their peers (Shaw et al. 2006). These individuals also demonstrate enhanced lateralization and more efficient interhemispheric communication, which are believed to be the basis for superior cognitive performance (Van Den Heuvel et al. 2009; Langer et al. 2012; Qi et al. 2019; Santarnecchi et al. 2015).

However, much of this research has focused on structural measures of the brain, leaving the functional aspects of these findings less well understood. The development of Homotopic Functional Affinity (HFA), a novel functional magnetic resonance imaging of the resting state (rfMRI), offers a promising method for examining interhemispheric interactions over time (Chen and Zuo 2024). Unlike traditional methods that focus on local correlations between homotopic regions, HFA incorporates global brain network organization, providing a more comprehensive view of how functional homotopy supports cognitive abilities. This study uses longitudinal data and the advanced HFA methodology to explore the development of functional homotopy from childhood to adolescence and its relationship with intelligence. By examining the dynamic changes in HFA over time, our aim is to uncover the neural mechanisms linking hemispheric coordination to cognitive performance. Specifically, we investigate how individuals with varying levels of intelligence differ in their development of functional homotopy. The findings of this research will not only deepen our understanding of the neural basis of intelligence, but also offer valuable information to tailor educational strategies and cognitive interventions to optimize individual learning and development.

## 2. Methods

### 2.1 Participants

Data were obtained from 179 participants (91 females, 88 males) aged 6.0 to 16.9 years as part of the developing Chinese Color Nest Project in Chongqing (devCCNP-CKG, Fan et al. 2023; Liu et al. 2021; Yang et al. 2017). The devCCNP-CKG employed an accelerated longitudinal design in which participants completed a baseline test followed by two follow-up evaluations at approximately 15-month intervals. In total, 351 visits were included in the current study: 64 participants completed one visit, 58 participants completed two visits, and 57 participants completed all three visits. Based on previous findings (Dong et al. 2021), data from all visits were divided into two groups: children (194 visits), adolescents (157 visits), using a cutoff age of 12 years to facilitate subsequent group-level analysis(Fig. 1).

**Fig. 1.**
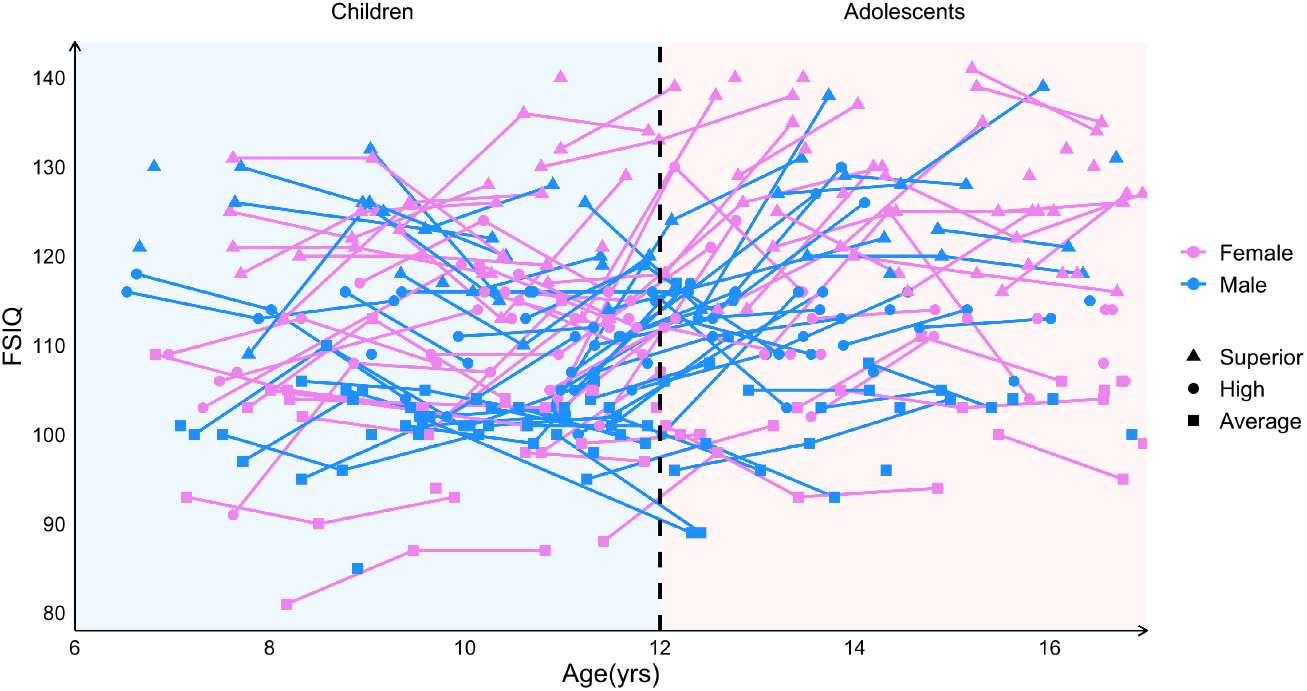
Longitudinal tracking of full-scale IQ (FSIQ) across age and gender. Each line connects data points from the same participant, illustrating individual FSIQ trajectories. Gender differences are highlighted by color coding—violet for females and blue for males. Participant IQ categories are represented by distinct shapes: triangles denote the superior group, circles the high group, and squares the average group. A vertical dashed line at age 12 categorizes participants into children and adolescents, facilitating developmental comparisons.

Data acquisition took place between March 2013 and January 2017. Participants were recruited from local primary and middle schools in Chongqing. All participants underwent comprehensive physiological and behavioral assessments. The inclusion criteria were the following: 1) no physical developmental abnormalities, 2) no history of psychological or psychiatric disorders, including family history, 3) the Child Behavior Checklist (CBCL, Achenbach and Edelbrock 1991) score below 70, 4) the Full Scale Intelligence Quotient (FSIQ) of the Chinese version of the Wechsler Intelligence Scale for Children, Fourth Edition (WISC-IV, Wechsler 2003) above 80, and 5) eligibility for MRI scan requirements. This project was approved by the Institutional Review Board of the Institute of Psychology of the Chinese Academy of Sciences. Informed consent was obtained from all participants’ guardians.

### 2.2. Intelligence assessment

Intelligence was assessed using the WISC-IV. Participants completed ten subtests: similarities, vocabulary, comprehension, digit span, letter-number sequencing, block design, picture concepts, matrix reasoning, coding, and symbol search. These subtests derive four distinct cognitive indexes: verbal comprehension, working memory, perceptual reasoning, and processing speed. Finally, these four indices are combined to generate the FSIQ, which reflects general intellectual ability.

### 2.3. MRI data acquisition

All MRI data were collected using a 3.0-T Siemens Trio MRI scanner. A 3D MPRAGE sequence was used to obtain an 8 minutes 19 seconds T1w MRI image (flip angle = 8.0^*°*^, TE = 3.02 ms, TR = 2600 ms, FOV = 256 mm, matrix = 256 × 256, voxel size = 1 mm × 1 mm × 1 mm). Before and after the T1w MRI scan, two rs-fMRI runs with identical parameters were performed using an echo planar imaging (EPI) sequence (flip angle = 80^*°*^, TE = 30 ms, TR = 2500 ms, FOV = 216 mm, matrix = 72 × 72, voxel size = 3 mm × 3 mm × 3.33 mm). Each rs-fMRI run consisted of 184 volumes, with 38 axial slices per volume, acquired over a total duration of 7 minutes and 45 seconds. During scanning, participants were instructed to remain motionless, awake, and fix their gaze on a “+” in the center of the screen.

### 2.4. MRI data preprocessing

Data preprocessing was performed by the Connectome Computation System (Xu et al. 2015), which integrates several publicly available toolboxes, including FreeSurfer (Fischl 2012), AFNI (Cox 1996), and FSL (Jenkinson et al. 2012). First, T1w images were skull-stripped using a modified pre-trained model based on Wang et al. (2021). The surface reconstruction and strict quality control were then performed to facilitate surface-based analyses. To compute HFA within a left-right symmetric space and enable comparison with adult HFA maps, the reconstructed surface was registered to the standard 32k grayordinate space using Ciftify (Dickie et al. 2019).

For rs-fMRI data, the first four volumes were removed to ensure signal stability. Then head motion correction, slice timing correction, and despiking were applied. The rs-fMRI images were then aligned with the T1w images using boundary-based registration (Greve and Fischl 2009). Potential artifacts from head motion, white matter, and cerebrospinal fluid signals were regressed using ICA-AROMA (Pruim et al. 2015). Subsequently, the rs-fMRI time series was resampled on the reconstructed 32k surface. Finally, a Gaussian smoothing kernel of 6 mm was applied to the time series data on the cortical surface for spatial smoothing. The Mean Framewise Displacement (mFD) was calculated for both rs-fMRI time series acquired during each visit. If either of the two time series from a visit exhibited an mFD greater than 0.5 mm, the data from that visit were excluded from the study.

### 2.5. HFA calculation

We calculated the HFA at the group and individual levels. To minimize the impact of random noise on rs-fMRI data and improve computational efficiency, the MELODIC’s Incremental Group Principal Component Analysis (MIGP, Smith et al. 2014) was applied to concatenate the time series. For the group-level analysis, the time series from all scans of all participants in the group were combined to create a group-level time series. For individual-level analysis, the time series from two rs-fMRI scans collected during a single visit were concatenated to generate a representative time series for that visit.

Given the symmetrical nature of grayordinate space, vertices with the identical label in the left and right hemispheres were considered homotopic. To calculate HFA, we first derived the functional connectivity profiles of the whole brain for each vertex. Next, we calculated the cosine similarity between the whole-brain connectivity profiles of each pair of homotopic vertices and applied Fisher’s z transformation to obtain the HFA. Notably, when cosine similarity between homotopic pairs was calculated, intrahemispheric connectivity sequences were aligned with intrahemispheric connectivity sequences, while interhemispheric connectivity sequences were aligned with inter-hemispheric connectivity sequences.

### 2.6. Development-HFA-Intelligence association analysis

The developmental effects of HFA were first examined by comparing the differences between the children and adolescent group maps. At the same time, differences in HFA at each cortical vertex across the two groups were statistically analyzed using Mann-Whitney U tests based on individual HFA maps.

Subsequently, Pearson’s correlations between FSIQ and HFA were calculated separately for children and adolescents to identify the relationship between HFA and intelligence and the possible developmental discrepancies.

Next, participants were classified into three IQ groups—superior, high, and average—based on their FSIQ scores, with each group representing one third of the total sample. For participants with repeated measures, their group classification was determined using the median FSIQ score across all of their visits. Group-level HFA maps were generated for six subgroups, defined by a combination of developmental stage (Children and Adolescents) and IQ classification. Among the 351 visits, the Adolescents-Average IQ subgroup had the smallest sample size (n = 41). To ensure comparability and minimize bias arising from unequal sample sizes, 40 visits were randomly selected from each subgroup to construct the final group-level HFA maps. In addition, Generalized Additive Mixed Models (GAMM) were applied to individual-level HFA maps to characterize developmental trajectories of whole-brain average HFA and vertex-wise HFA values across IQ groups. The models were constructed using the mgcv package in R, as shown in equation (1).

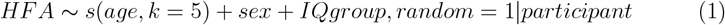

The *s*() represents a penalized smoothing spline function of age to fit *HFA, k* denotes the degrees of freedom, *sex* is the fixed effect to control potential sex effects, *IQgroup* is the fixed effect to account for IQ differences, *random* represents the random intercept for each participant, accounting for the repeated measures across visits.

Using established models, HFA was predicted at each vertex at successive ages of 6.0 to 16.9 years. The Coefficient of Variation (CV) for the predicted HFA at each vertex was subsequently calculated to evaluate the developmental effects of HFA (2).

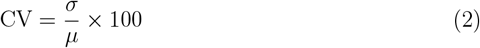

*σ* denotes the standard deviation of the model-fitted HFA for a given vertex over the age range of 6.0 to 16.9 years, while *µ* represents the mean of the predicted values.

## 3. Results

### 3.1. Age-related decrease in HFA

The HFA group maps of children and adolescents revealed significant effects related to age. The HFA map of the children’s group showed an early emerging adult-like pattern (Chen and Zuo 2024), while the map of the adolescent group showed a more pronounced adult-like HFA pattern (Fig. 2A). In general, children showed higher HFA, while adolescents demonstrated lower global HFA. The most pronounced decline in HFA in adolescents was observed in the Ventral Attention Network (VAN), the Fronto-Parietal Network (FPN), and the Default Mode Network (DMN). Vertex-level Mann-Whitney U tests revealed consistent developmental patterns along the unimodal-to-transmodal gradient. The primary cortices exhibited the weakest developmental effect sizes, while the associative cortices exhibited the strongest developmental effect sizes (Fig. 2B).

**Fig. 2.**
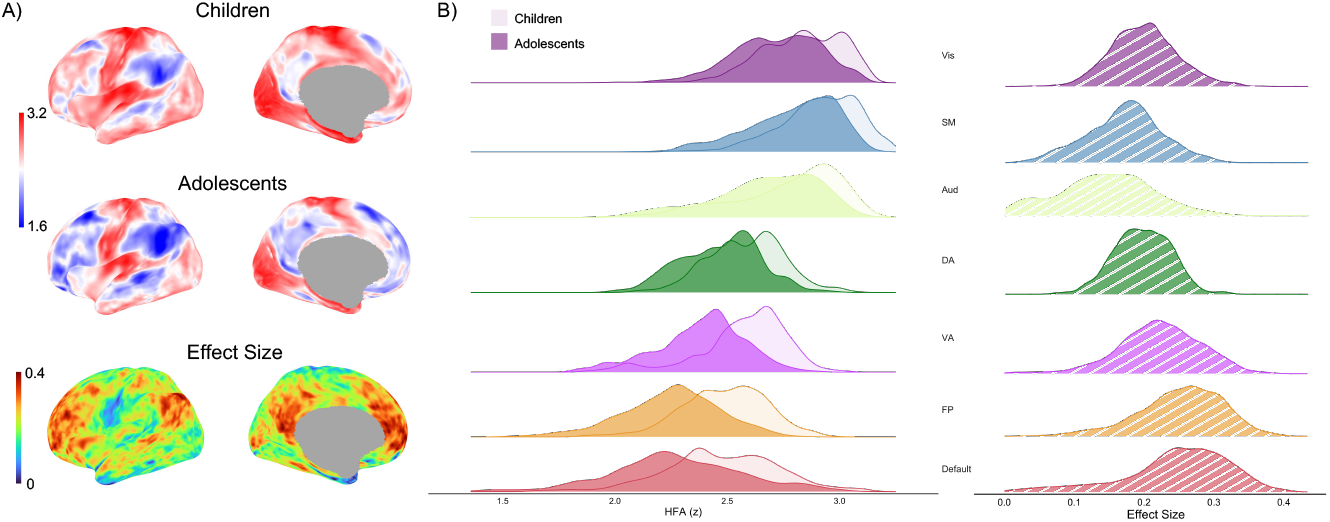
HFA Maps and their comparative analyses between children and adolescents. A) Whole-brain HFA maps for children and adolescents, accompanied by an effect size map from Whitney U tests that highlight significant developmental differences. B) HFA distributions across functional networks, as defined by the parcellation scheme from the Chinese Human Connectome Project (Ge et al. 2023), developed using the methodology described in Kong et al. (2019). Lighter shades (higher transparency) represent children, while darker shades denote adolescents. Comparative effect sizes of inter-group differences, detailed for each network, are presented in the last column.

### 3.2. Enhanced negative HFA-FSIQ association in adolescents

In the children group, the correlation coefficients between HFA and FSIQ fell within a 95% confidence interval ranging from -0.14 to 0.13. The regions exhibiting the strongest negative correlations were located primarily in the FPN and VAN, particularly in the prefrontal cortex and insular regions. In contrast, the regions with the strongest positive correlations were detected mainly within the somatosensory cortex and the dispersed areas of the DMN (Fig. 3).

**Fig. 3.**
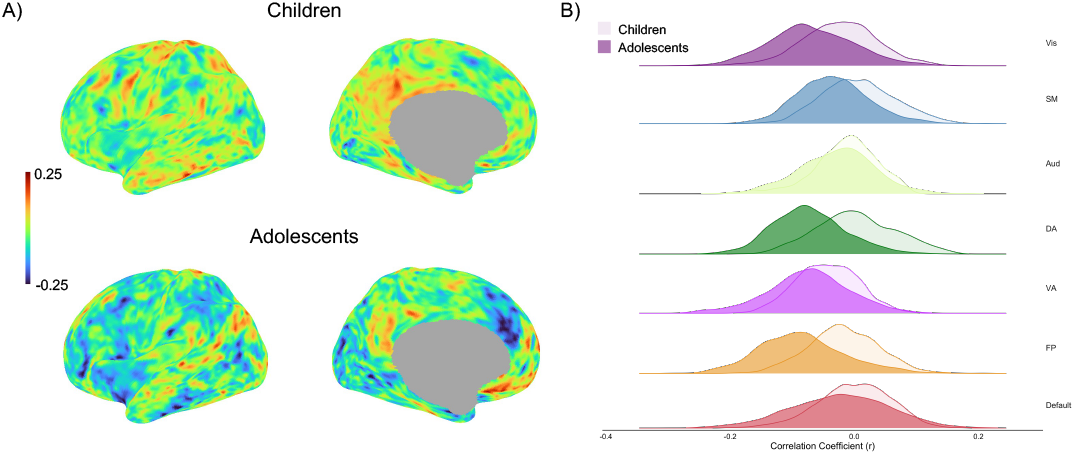
Correlation maps of HFA and FSIQ in children and adolescents. A) Correlation coefficient maps at the vertex level depicting the relationship between HFA and FSIQ for both children and adolescent groups. B) Distribution of correlation coefficients across different brain networks, illustrating the regional and group variability in the relationship between HFA and cognitive performance. Lighter shades (higher transparency) represent children, while darker shades denote adolescents.

Compared to children, adolescents showed a stronger overall negative correlation. In this group, 95% of the correlation coefficients of the vertices range from -0.2 to 0.14. The brain regions that exhibited a strong negative correlation in the children’s group demonstrated a more pronounced negative correlation in adolescents. In addition, a wider range of prefrontal areas exhibited negative correlations, along with extensive regions within the DMN. In particular, the strongest positive correlations were located in the subregions of the medial prefrontal cortex, the inferior parietal lobule, and the posterior cingulate cortex, all of which are recognized as components of DMN (Fig. 3). However, we note that, after applying the False Discovery Rate (FDR) correction, only a few isolated vertices showed significant correlations in both groups.

#### 3.3. Developmental rate differentiation across IQ groups

The basic demographic characteristics for different IQ groups are presented in Fig.1. In the Children group maps, subtle differences were observed in HFA distributions across IQ groups, with the superior IQ group exhibiting slightly lower HFA values. In contrast, within the Adolescents group, disparities in HFA maps between IQ groups became more pronounced, particularly with the superior IQ group showing the lowest HFA values (Fig. 4A). These patterns were further supported by GAMM analyses of individual-level whole-brain average HFA, which revealed that the superior IQ group exhibited the fastest global rate of HFA development. In comparison, the high-IQ and average-IQ groups demonstrated slower, yet similar, HFA decrease rates. By age 17, the superior IQ group showed the lowest HFA values, whereas the average-IQ and high-IQ groups displayed comparably higher HFA levels (Fig.4B).

**Fig. 4.**
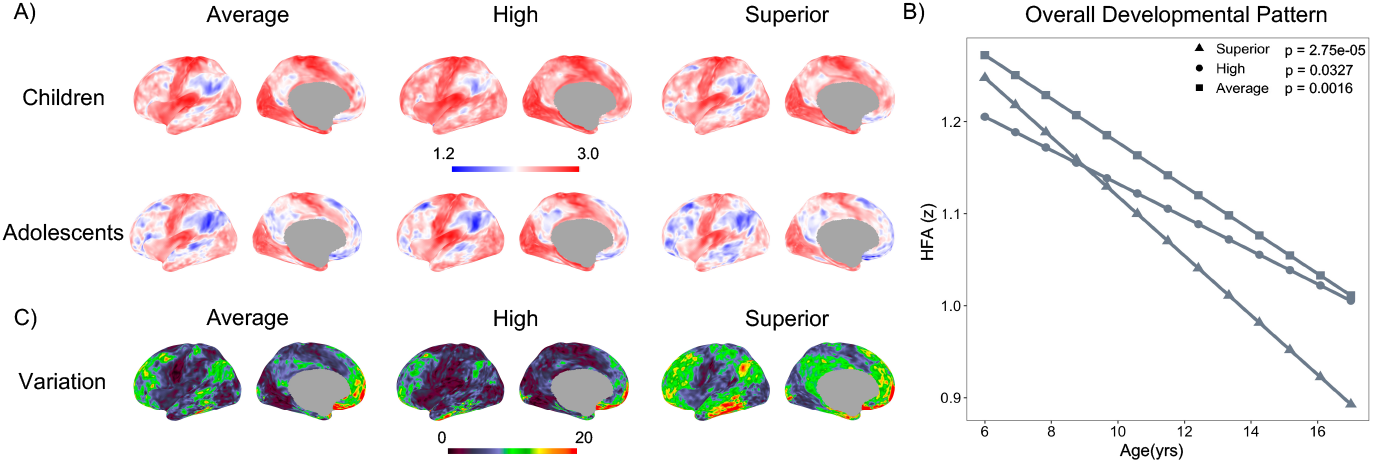
IQ-dependent differences in HFA development. A) Group-level HFA maps across IQ groups during childhood and adolescence. B) Developmental trajectories of whole-brain average HFA of each IQ groups. Each group are represented by distinct shapes: triangles denote the superior group, circles the high group, and squares the average group. C) Vertex-wise coefficient of variation (CV) maps of HFA developmental trajectories across IQ groups.

The vertex-wise CV maps derived from developmental modeling closely mirrored the patterns observed in group maps and whole-brain mean value analyses (Fig.4C). The superior IQ group demonstrated the most widespread developmental changes, characterized by higher CV values across multiple functional networks compared to the high-IQ and average-IQ groups. Specifically, elevated CV values in the superior IQ group were observed in both higher-order networks and primary networks, including the DMN, FPN, visual, auditory, somatosensory, and dorsal attention networks (DAN). In contrast, the high-IQ and average-IQ groups primarily showed significant developmental effects within higher-order networks, while age-related changes within primary networks were minimal. Moreover, the spatial distribution of CV maps across IQ groups was negatively correlated with adult HFA maps. Regions with high HFA values in adulthood exhibited limited developmental change, whereas regions with lower adult HFA values showed more pronounced age-related trajectories.

## 4. Discussion

This study provides a comprehensive examination of the developmental trajectory of functional homotopy and its association with intelligence. The findings reveal that the functional specialization of the hemispheric region increases progressively during development, following a gradient from unimodal to transmodal cortices. IQ-superior individuals demonstrate a more rapid reduction in functional homotopy, reflecting advanced hemispheric specialization and neural efficiency. These results highlight the intricate interplay between brain maturation and cognitive development, emphasizing the importance of dynamic interhemispheric coordination in shaping individual cognitive abilities.

### 4.1. Consistent spatial and temporal patterns of functional homotopy along the unimodal-transmodal axis

Previous research has provided substantial evidence for an age-related increase in hemispheric specialization. Specifically, the mean homotopic functional connectivity across the whole brain gradually decreases in early development (Zuo et al. 2010), regions associated with language skills (e.g. the prefrontal cortex) or visuospatial skills (e.g. the temporal and parietal lobes) exhibit increased lateralization with age (Qi et al. 2019; Gracia-Tabuenca et al. 2018; Cai et al. 2019). Despite these findings, a comprehensive understanding of the patterns and developmental trajectories of functional homotopy in all brain regions remains elusive. This study addressed this gap by revealing the function of homotopic regions specialized spatially and temporally congruent with the unimodal-transmodal axis gradient.

The unimodal-transmodal gradient has emerged as a fundamental framework for understanding the evolutionary, developmental, and genetic underpinnings of brain structure and function. This gradient is widespread in different aspects of the human brain (Huntenburg et al. 2018; Sydnor et al. 2021). Along this gradient axis, the unimodal cortices, which are primarily responsible for basic sensory perception, emerged earlier in evolution and typically matured earlier in development. In contrast, maturation of later evolving transmodal cortices requires a longer time and is related to the emergence of advanced cognitive functions (Buckner and Krienen 2013; Hill et al. 2010). Our findings further reinforce the critical role of this gradient in shaping the intrinsic functional organization of the human brain, particularly through the lens of hemispheric functional integration and specialization throughout development.

Functional homotopy appears to reflect the overall functional organization in the human brain (Labache et al. 2023). Generally, unimodal cortices tend to engage in intense bilateral dialogue, whereas transmodal regions prioritize intrahemispheric interactions (Wang et al. 2014). In unimodal cortices, elevated levels of functional homotopy contribute to maintaining interhemispheric sensory coherence and motor coordination, allowing rapid and consistent responses to external stimuli. In contrast, in transmodal cortices, the increase in interhemispheric specialization supports the demands of higher-order cognitive processes, thereby enhancing the efficiency and flexibility of information processing in the brain (Toga and Thompson 2003; Gee et al. 2011). Meanwhile, the differential developmental trajectories of functional homotopy along the unimodal-transmodal cortices reflect the natural progression of the brain from basic perceptual abilities to complex cognitive functions. Throughout brain development, the configuration of the human connectome progressively changes from a “local” to a more “distributed” pattern (Fair et al. 2009). This transformation involves a transition from distance-driven constraints to those shaped by topological properties (Zuo et al. 2017), enabling functional networks to become increasingly differentiated, which may mediate the development of cognition during adolescence (Gu et al. 2015; Baum et al. 2017; Uddin et al. 2011). This process is particularly pronounced in the transmodal cortices, where intrinsic functional differentiation and pluripotency are activated during late childhood. These changes are evidenced by an increase in the diversity of interactions between transmodal cortices and other brain regions, along with increased complexity in their activity patterns when responding to cognitively demanding tasks (Gu et al. 2015). As these transformations unfold, the functional variability in homotopic brain regions was amplified, leading to a reduction in HFA values.

The consistency between spatial distribution and developmental patterns not only promotes the efficient functioning of sensory and motor systems but also facilitates the optimization of advanced cognitive processes, ultimately achieving an effective balance between coordination and cognitive adaptability.

### 4.2. The dynamic nature of brain-mind association

The evolving relationship between HFA and IQ in different age groups highlights the dynamic nature of brain-mind association, consistent with findings reported in broader research on human brain development (Shaw et al. 2006; Shaw 2007; Li et al. 2024; İçer 2020). In the current study, age-related correlation changes were particularly pronounced in most within the VAN, FPN, and DMN.

Recent findings suggest that VAN maturation plays a crucial role in facilitating age-dependent changes in cortical organization and cognitive function (Dong et al. 2024). This network establishes an intricate structure, particularly the thalamus and amygdala, both of which undergo significant developmental changes during puberty. Since these connections are predominantly ipsilateral, they may support hemispheric functional specialization in emotional and cognitive processing (Ghaziri et al. 2018; Duerden et al. 2013).

The FPN is a typical lateralized network that supports critical cognitive functions such as language processing and executive control, both of which exhibit marked hemispheric specialization (Wang et al. 2014). During adolescence, the development of the frontoparietal cortices mediates the maturation of executive functions (Baum et al. 2017). After adolescence, the cortical thickness asymmetry in the frontal and parietal lobes increases progressively (Zhou et al. 2013). Schlaggar et al. (2002) reported that an increase in leftward asymmetry in language-related regions was associated with the growth of linguistic skills. In the context of language tasks, children demonstrate bilateral activation of the inferior frontal cortex, whereas adults predominantly exhibit left-hemispheric activation (Olulade et al. 2020).

The changes observed within the DMN are particularly intriguing. During childhood, correlations within the DMN are largely positive; however, during adolescence, while some regions maintain strong positive correlations, others develop pronounced negative correlations. This shift signifies a differentiation in brain-mind relationships within the DMN, suggesting a more nuanced organization as cognitive demands evolve. Previous studies have demonstrated that the DMN comprises at least two spatially juxtaposed subsystems, each associated with distinct cognitive domains (Braga et al. 2019). The changes documented in this study may reflect the differential cognitive associations of these DMN subsystems and highlight the role of hemispheric specialization in shaping their functional contributions.

### Individual differences in intelligence reflected by brain growth

Although the direct correlation between HFA and intelligence observed in this study was relatively weak, comparisons of correlation patterns between age groups and HFA developmental trajectories in different IQ groups revealed significant divergence in the developmental course of functional homotopy between these groups. This divergence is consistent with the findings of previous research on the relationship between brain morphology and intelligence (Shaw et al. 2006; Schnack et al. 2015). These results suggest that intelligence may be more closely linked to the dynamic features of brain development, such as the magnitude and timing of developmental changes, rather than to static brain measures per se. Therefore, researchers must prioritize the study of the dynamic processes that underlie brain maturation when exploring the neural basis of intelligence.

The age-related development patterns of HFA in different IQ groups consistently follow the unimodal-to-transmodal axis. However, the superior IQ group exhibits the fastest rate of HFA decline across the entire brain, with a nearly linear decrease observed between 6.0 and 16.9 years of age. This rapid decrease in HFA suggests an accelerated development of hemispheric functional specialization, where the ability of the left and right hemispheres to independently process information and engage in specialized functions is significantly enhanced. Through this process, homotopic regions become more efficient in their division of labor and coordination, thus supporting the integration and processing of information required for complex cognitive tasks. This developmental trajectory may provide a foundation for the superior performance of individuals with a higher IQ in future complex cognitive challenges. Studies on brain-intelligence relationships in adults lend some support to this inference. Research indicates that adults with higher IQ typically demonstrate greater global and local network efficiency, which supports more effective parallel information processing (Van Den Heuvel et al. 2009; Li et al. 2009; Langer et al. 2012). In addition, better cognitive performance is associated with increased network flexibility during task switching, which enables faster reconfiguration of connections between functional networks (Hearne et al. 2017).

Only the superior IQ group showed a marked age-related decline in HFA within primary functional networks. Established research has consistently indicated that people with higher intelligence exhibit faster reaction times and greater efficiency in processing visual and auditory stimuli compared to those with lower intelligence (Deary et al. 2010). In addition, studies have reported lower homotopic connectivity in the primary cortices of highly intelligent individuals, including regions such as the visual cortex, the somatosensory cortex, and the supplementary motor area (Santarnecchi et al. 2015). When processing basic sensory information, the overwhelming influx of sensory input may exceed the brain’s capacity to effectively manage and process the information. Therefore, the ability to suppress irrelevant stimuli becomes a key determinant of perceptual efficiency. Enhanced hemispheric specialization in primary sensory cortices is likely to facilitate improved perceptual efficiency and information processing capabilities, which in turn supports more advanced cognitive functions. This mechanism may provide an explanation for the significant development of hemispheric specialization observed in the superior IQ group within the primary cortex.

## 5. Conclusion

This study highlights the developmental trajectory of functional homotopy and its relationship with intelligence, offering novel insights into the neural mechanisms underlying cognitive development. Using the innovative HFA methodology, we demonstrate that functional homotopy decreases with age, particularly in higher-order networks, and is closely associated with individual differences in intelligence. These findings improve our understanding of typical brain development and provide a foundation for educational and cognitive interventions tailored to individual needs.

Future research should include the analysis of subcortical structures, which are critical to brain connectivity and cognitive function. Expanding HFA to capture interactions between the cortical and subcortical regions could provide a more comprehensive view of brain development. In addition, HFA shows great potential for basic research and clinical applications, such as frequency-dependent information on dark brain energy (Gong and Zuo 2025), early diagnosis, and the study of neurodevelopmental disorders. By identifying functional homotopy abnormalities, this method could offer new tools to understand the mechanisms behind cognitive impairments and developmental disorders, paving the way for improved diagnostic and therapeutic strategies.

## Declaration of interests

The authors declare that they have no known competing financial interests or personal relationships that could have appeared to influence the work reported in this manuscript.

## Data availability

The datasets analyzed in this study are publicly available from Chinese Color Nest Project - Lifespan Brain-Mind Development Data Community (https://ccnp.scidb.cn/en).

## Acknowledgment

The STI 2030 - the major projects of the Brain Science and Brain-Inspired Intelligence Technology (2021ZD0200500), the Start-up Funds for Leading Talents at Beijing Normal University, the National Natural Science Foundation of China (82102134, 82322035, 62273076), the Key-Area Research and Development Program of Guangdong Province (2019B030335001), the Beijing Municipal Science and Technology Commission (Z161100002616023, Z181100001518003), the Major Project of National Social Science Foundation of China (20&ZD296), the CAS-NWO Programme (153111KYSB20160020), the Chinese Academy of Sciences Key Research Program (KSZD-EW-TZ-002), the Major Fund for International Collaboration of National Natural Science Foundation of China (81220108014) and the National Basic Research (973) Program (2015CB351702) provide funding supports.

## Notes

https://ccnp.scidb.cn/en

### Competing Interest Statement

The authors have declared no competing interest.

